# Genetic architecture of oxidative stress tolerance in the fungal wheat pathogen *Zymoseptoria tritici*

**DOI:** 10.1101/2020.02.20.957431

**Authors:** Ziming Zhong, Bruce A. McDonald, Javier Palma-Guerrero

## Abstract

Reactive oxygen species are toxic byproducts of aerobic respiration produced during cell growth. They also are an important component of plant defenses to inhibit microbial pathogens. Tolerance to oxidative stress contributes to viability and pathogenicity of plant pathogens. However, the complex molecular network of oxidative stress responses hinders identification of the genes contributing to variation in this trait. Variation in genes affecting responses to oxidative stress is likely to affect the evolutionary potential of pathogen tolerance to host defences. Here, we employed a forward genetic approach to investigate the genetic architecture of oxidative stress tolerance in the fungal wheat pathogen *Zymoseptoria tritici*. By performing quantitative trait locus (QTL) mapping in two crosses, we identified several genomic regions associated with tolerance to oxidative stress, including a QTL having a large effect on growth under oxidative stress. We found evidence for a significant trade-off between growth under non-stressful conditions and growth inhibition under oxidative stress. We identified a large QTL associated with this trade-off and with growth under non-stressful conditions, suggesting that differences in fungal growth could result in different sensitivities to oxidative stress. Our results suggest that genes related to fungal growth could also contribute to variation in oxidative stress tolerance among fungal strains.

## INTRODUCTION

Reactive oxygen species (ROS) are toxic free radicals that can oxidize biomolecules and disturb basic metabolism. Aerobic microbes produce a large amount of ROS that may interfere with metabolism as a normal byproduct of respiration during cell growth (Heller & Tudzynski 2011; Segal & Wilson 2018). Cells can also be exposed to an endogenous ROS burst when experiencing environmental changes and abiotic stresses, such as thermal stress (Trachootham *et al*. 2008; Gocheva *et al*. 2009; Heller & Tudzynski 2011). During plant-pathogen interactions, ROS are secreted by plants within minutes to act as secondary signals to trigger plant defenses, and then peak during the hypersensitive response to isolate pathogens locally (Doehlemann & Hemetsberger 2013). Hence, the efficient elimination of excessive ROS and the ability to grow under oxidative stress are important traits for fungal pathogens during plant infection.

Different chemicals and enzymes can be synthesized to counteract the effects of oxidative stress (Thorpe *et al*. 2004; Heller & Tudzynski 2011; Segal & Wilson 2018). Enzymes such as superoxide dismutases (SOD), catalases, and catalase-peroxidases are induced in oxidative stress environments (Broxton & Culotta 2016; Papadakis & Workman 2014). Reductases and peroxidases employing thioredoxin and glutathione as electron donors were also reported to be important for ROS reduction (Fernandez & Wilson 2014; Yang *et al*. 2016; Ma *et al*. 2018). The NADPH oxidase complex (NOX) involved in ROS generation was also reported to be associated with sensitivity to oxidative stress (Chen *et al*. 2014). Other molecules found in fungi, including melanin and carotene, were also suggested to protect cells from ROS (Jacobson 2000; Heller & Tudzynski 2011). Oxidative stress responses were found to be largely regulated by the transcription factor Yap1 and the MAPK pathways (Molina & Kahmann 2007; Segmüller *et al*. 2007; Walia & Calderone 2008; Lin *et al*. 2009; Ronen *et al*. 2013; Yu *et al*. 2017; Segal & Wilson 2018). Genes that regulate many stress responses in fungi, referred to as core environmental stress response (CESR) genes, such as Msn2 and Msn4, were also found to be involved in the oxidative stress response in fission yeast (Chen *et al*. 2003; 2008).

Results from previous studies suggested that other unknown genes also affect oxidative stress tolerance and that there are genes with dispensable and overlapping functions that decrease the effects of ROS (Grant *et al*. 1997; Rolke *et al*. 2004; Gohari *et al*. 2015; Segal & Wilson 2018). Transcriptomic studies in fission yeast showed that a complex network of hundreds of genes is involved in the oxidative stress response (Chen *et al*. 2008). This complex network was affected by the exposure time, ROS dose, ROS location and the species. In such complex gene networks, it is difficult to determine which genes contribute to the tolerance to oxidative stress and whether these genes retain a high evolutionary potential in natural populations. Most previous studies in fungal plant pathogens determined the functions of major ROS scavengers by analyzing single gene mutations (Heller & Tudzynski 2011; Segal & Wilson 2018). We hypothesized that association mapping would provide more powerful insights into the evolution of this complex trait by enabling a whole-genome perspective. Combining genotypic and phenotypic variation can enable identification of important genomic regions or genes that contribute to the variation in oxidative stress tolerance. Association mapping studies utilizing wild-type strains can provide insight into the evolution of this trait in natural populations.

The fungus *Zymoseptoria tritici* has been successfully used in quantitative trait locus (QTL) mapping and genome-wide association studies (GWAS) (Lendenmann *et al*. 2014; 2015; 2016; Hartmann *et al*. 2017; Stewart *et al*. 2018). This pathogen reduces wheat yields globally by damaging infected leaves (Torriani *et al*. 2015). *Z. tritici* is a latent necrotroph with a long asymptomatic phase that varies according to the combination of pathogen strain and host cultivars (Sanchez-Vallet *et al*. 2015). Previous studies in *Z. tritici* considered the importance of ROS during plant infection. Two bursts of hydrogen peroxide were detected during a compatible infection: a weak burst during the latent phase (around 5 days post inoculation, dpi) and a strong burst that occurred during the start of the necrotrophic phase (around 13 dpi) (Shetty *et al*. 2003). It was found that infiltration of catalase into infected leaves increased disease symptoms and accelerated disease development, while infiltration of hydrogen peroxide (H_2_O_2_) decreased the disease symptoms and delayed fungal sporulation (Shetty *et al*. 2007). These experiments illustrated the importance of ROS tolerance during plant infection.

Some genes involved in the oxidative stress response have already been characterized in *Z. tritici*. A catalase-peroxidase (*ZtCpx1*) was found to be essential for growth *in vitro* in the presence of hydrogen peroxide and for disease development during plant infection (Gohari *et al*. 2015). Another catalase-peroxidase (*ZtCpx2*) was found to be dispensable for the *in vitro* oxidative stress response but it was important for the transition from the biotrophic to the necrotrophic phase during plant infection (Gohari *et al*. 2015). The major oxidative stress regulator Yap1 was also identified in *Z. tritici*, and was found to be essential for *in vitro* tolerance to oxidative stress but not for *in planta* virulence (Yang *et al*. 2015). These results confirmed the importance of some known ROS management systems in *Z. tritici* and illustrated the need for further studies to identify other genes involved in oxidative stress tolerance.

In this study, we investigated the genetic architecture of oxidative stress tolerance in *Z*. *tritici* by performing quantitative trait locus (QTL) mapping of tolerance to hydrogen peroxide under axenic conditions in two segregating populations. Significant QTLs were compared in the presence or absence of oxidative stress to infer the biological functions of the QTLs. Large-effect QTLs associated with tolerance to oxidative stress were further investigated to identify genes related to oxidative stress tolerance by combining previously published *in planta* and *in vitro* RNAseq datasets. We identified several QTLs associated with oxidative stress tolerance, including one QTL strongly associated with growth under oxidative stress. Our data also indicated a trade-off between growth under non-stressful conditions and growth inhibition under oxidative stress, indicating that genes contributing to general growth may also affect stress tolerance. This study suggests that the natural variation in oxidative stress tolerance could reflect variation both in genes specific for oxidative stress response and genes related to fungal growth under non-stressful conditions.

## MATERIALS AND METHODS

### Phenotyping for oxidative stress tolerance in vitro

Two crosses among four Swiss strains of *Z. tritici* (1E4 × 1A5 and 3D1 × 3D7) generated 249 and 257 offspring, respectively, as reported in previous studies (Lendenmann *et al*. 2014; 2015; 2016). The four parents and their 506 progeny were separated into 20 sets to phenotype for oxidative stress tolerance as follows. Spores were recovered from glycerol stocks kept at –80 °C and grown in YPD (yeast potato dextrose) media for 5 days at 18 °C on a shaker at 120 rpm. The spore suspensions were filtered through a double layer of sterilized cheesecloth and then centrifuged at 3273 *g* for 15 min. The spore pellets were diluted to a concentration of 200 spores per ml, and 200 μl of the spore solution was added to each PDA (Difco potato dextrose agar) Petri plate to obtain around 40 colonies per plate. To create the oxidative stress environment, 113 μl of 30 % H_2_O_2_ (SIGMA-ALDRICH Chemio GmbH) was added into one liter of autoclaved and cooled (< 60 °C) PDA media to produce a final concentration of 1.0 mM H_2_O_2_. Unamended PDA was used as the control environment. Each strain was inoculated onto three replicate Petri plates for each environment. All plates were incubated at 18 °C in the dark. Digital images of each plate were acquired at 8- and 12-days post inoculation (dpi) as described by Lendenmann *et al*. (2014).

To measure colony area (cm^2^) and the degree of melanization, all images were processed in MatLab R2017b (The MathWorks, Inc., Natick, Massachusetts, United States) using a purpose-developed script (File S1) based on instructions given in the Image Segmentation Tutorial by Image Analyst (https://ch.mathworks.com/matlabcentral/fileexchange/25157-image-segmentation-tutorial). Colony radii were calculated as 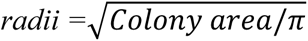. To measure melanization of each colony, gray scale values were obtained from RGB color values by using the *rgb2gray* function in MatLab. Higher gray values indicate lower melanization on the 0-255 gray scale, where 0 is completely black and 255 is completely white. The average colony radius and average gray value for each isolate under each condition was calculated for each plate, and the mean value across the three replicates was used as the phenotype input for growth and melanization in the QTL mapping. We also calculated the growth rate and melanization rate between 8 and 12 dpi, as the colony growth between these time points was found to be linear in a previous study (Lendenmann *et al*., 2015). As most strains showed different growth patterns under stressed and non-stressed conditions, the relative values of growth or melanization, representing the percentage of growth reduction or increase in melanization due to oxidative stress, were calculated for each strain to estimate differences in sensitivity to oxidative stress among offspring. All phenotypes used as inputs for QTL mapping were calculated as shown below, and all statistical analyses were performed in R (R Core Team, 2018).

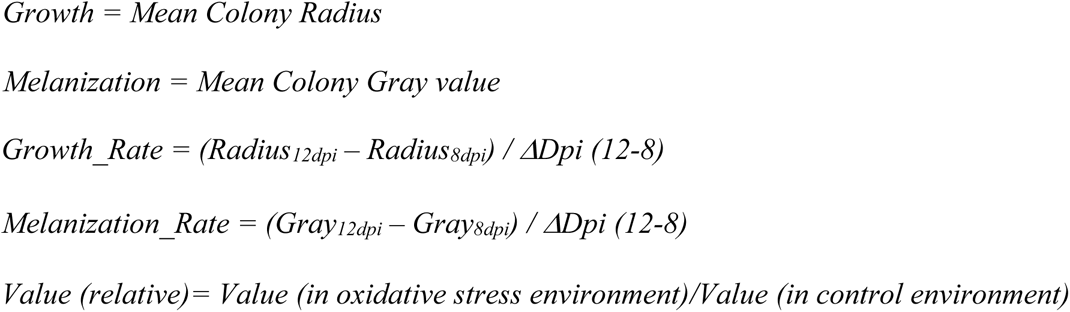

To estimate the phenotypic variation resulting from the genetic variation, broad-sense heritability was calculated as 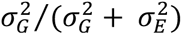 (Bloom *et al*. 2013), where 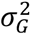 is the genetic variance and 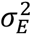 is the residual variance. The calculation of these two variance components was performed using mixed models with strain, replicate and set as random effects in the *sommer* package (Covarrubias-Pazaran 2016) in R. To detect transgressive segregation, we performed Dunnett’s test to identify any segregants with mean values either significantly higher or lower than the parents (p<0.001) (Johansen-Morris *et al*. 2006). The Dunnett’s tests were performed by using replicates in the *multcomp* package (Hothorn *et al*. 2008) in R.

### Genotype data and QTL mapping

SNP data used for QTL mapping were obtained from RADseq data generated previously (Lendenmann *et al*. 2014), but using the finished genome sequence of one of the parental strains (Plissonneau *et al*. 2018) as the reference genome for each cross (1A5 for the 1E4 × 1A5 cross and 3D7 for the 3D1 × 3D7 cross). The SNP markers were generated and filtered as described in previous studies (Zhong *et al*. 2017; Meile *et al*. 2018). This provided 35030 SNP markers in the 1E4 × 1A5 cross and 57513 SNP markers in the 3D1 × 3D7 cross, leading to average marker distances of 0.31 and 0.10 cM (1145 and 658 bp) respectively. Single-QTL genome scans using standard interval mapping were performed to provide an overview of the genetic architecture of each trait. All QTL analyses were performed using the *R/qtl* package (Broman *et al*. 2003) in R, following the instructions in ‘A Guide to QTL Mapping with R/qtl’ (Broman & Sen 2009). The significance of each QTL was assessed using 1000 permutation tests across the entire genome. Genes located within the 95% Bayes credibility intervals were identified according to the genome annotations of the reference parental strains (Plissonneau *et al*. 2018).

### Identification of candidate genes within QTL confidence intervals

Only QTLs with high LOD scores (LOD > 10) were included in the analyses aiming to identify candidate genes associated with each trait. All candidate genes were BLASTed to the NCBI database (https://blast.ncbi.nlm.nih.gov/Blast.cgi) to confirm their functional domains and to search for homology to functionally characterized genes. Sequence variation among the parental strains in the QTL confidence intervals was identified by performing sequence alignments (CLC Sequence Viewer 7.6.1, QIAGEN Aarhus A/S). The previously published *in planta* expression data (Palma-Guerrero *et al*. 2017) at 7 dpi and *in vitro* expression data (Francisco *et al*. 2019) obtained for the four parental strains were used to confirm the gene models and to compare the expression of candidate genes. Genes with at least one sequence variant among the parental strains in either the protein-encoding sequence or the 5’- or 3’- UTR regions were considered to be top candidates to explain the observed marker-trait associations.

### In planta expression of candidate genes

To clarify the possible *in planta* function of candidate genes, we used the *in planta* expression data obtained in a previous study for the 3D7 parent (Palma-Guerrero *et al*. 2016), which consisted of five time points, covering the complete life cycle of the fungus in wheat leaves. We analyzed the expression of the top candidate genes and compared them to genes known to encode domains involved in the oxidative stress response. For this analysis we focused only on genes with homology to genes already known to be related to oxidative stress in fungi.

## RESULTS

### Variation in growth and melanization

Two segregating F1 populations of *Z. tritici* were scored for growth and melanization in the presence and absence of hydrogen peroxide. Growth of nearly all strains was inhibited (i.e. relative growth was < 1) by 1 mM H_2_O_2_ at 8 dpi in both crosses (Figure 1A, left panel), but the degree of inhibition varied among strains. The 1E4 and 1A5 parental strains showed similar levels of growth inhibition (1A5 = 0.68 ± 0.01, 1E4 = 0.62 ± 0.04) under oxidative stress, but a total of 94 of their offspring exhibited growth inhibition significantly (p < 0.001) higher than 1A5 or lower than 1E4 (Figures 1A and 1B). We interpreted this as evidence of transgressive segregation. Parental strains 3D1 and 3D7 showed larger differences (3D1 = 0.81 ± 0.03, 3D7 = 0.52 ± 0.02) in the degree of growth inhibition, and their offspring exhibited a wider range in relative growth (0.44-1.13) than the parental strains (Figure 1A and 1B). Only eight offspring in this progeny population had significantly (p < 0.001) higher growth inhibition than 3D1, and no offspring had significantly lower growth inhibition than 3D7. At 8 dpi, only one offspring from the 1E4 × 1A5 cross and five offspring from the 3D1 × 3D7 cross did not show growth inhibition under oxidative stress. At 12 dpi, 25 offspring from the 1E4 × 1A5 cross and 47 offspring from the 3D1 × 3D7 cross showed an increase in growth under oxidative stress compared to the control (i.e. relative growth was > 1), suggesting a capacity in these offspring to negate the effects of oxidative stress. Weak but significant correlations between growth under control and oxidative stress conditions were found in both segregating populations (Figure 1C), suggesting inherent differences in growth rates among strains irrespective of their environment. Growth in the control environment and growth inhibition under oxidative stress were negatively correlated in both segregating populations (Figure 1D) at both time points, indicating that the strains that grew the fastest in the control environment were the most affected by oxidative stress.

**Figure 1.**
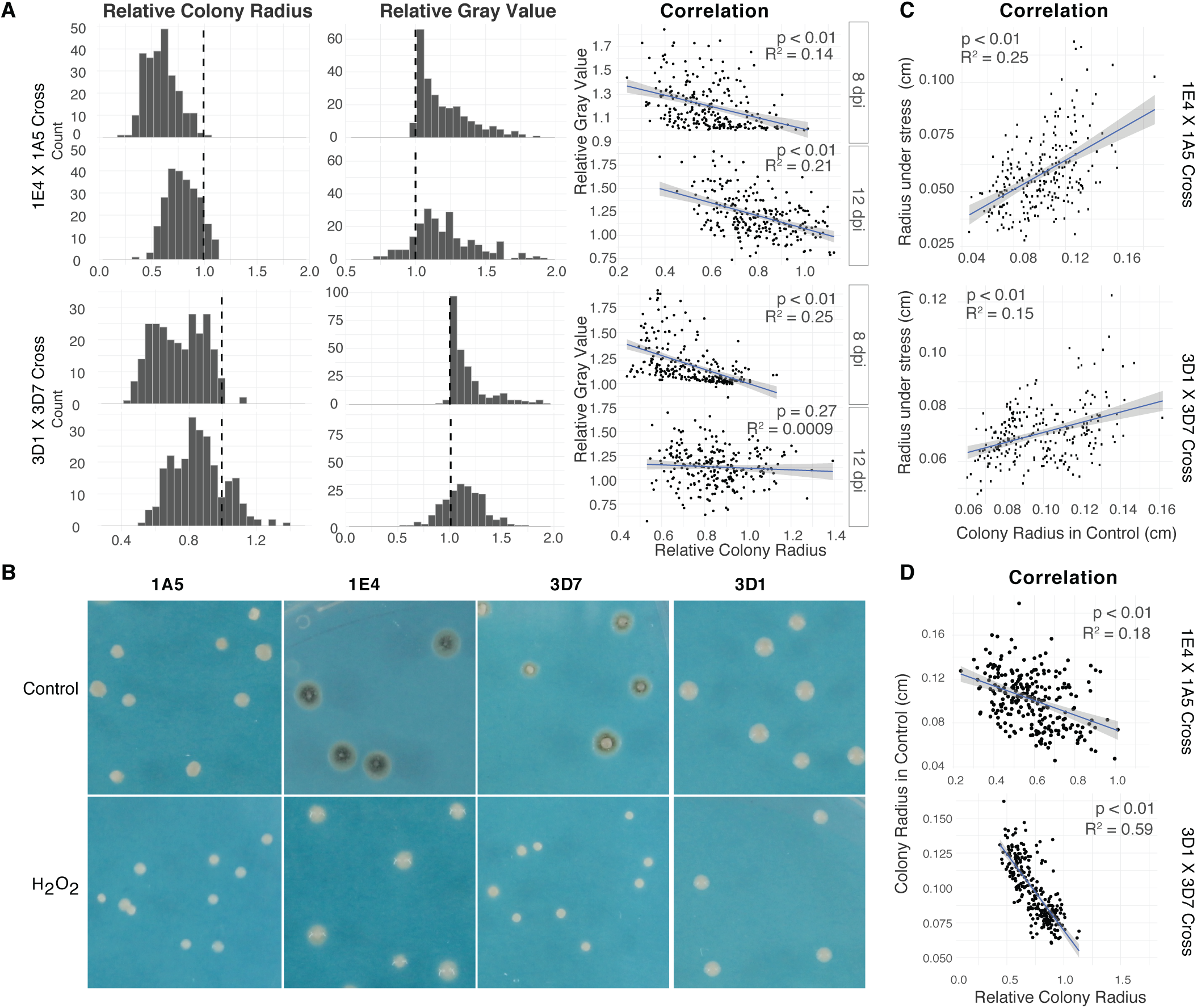
Variation in growth and in melanization under oxidative stress. (A) The distribution of the relative growth (left panels) and relative melanization (middle panels) traits in the 1E4 × 1A5 cross and the 3D1 × 3D7 cross. Relative colony radius (x axis) less than 1 indicates that oxidative stress inhibited the growth, and vice versa. Relative melanization value (x axis) less than 1 indicates more melanization under oxidative stress. The dashed line stands for 1, indicating that the value under oxidative condition equals the value under the control condition. The linear correlations between relative growth and relative melanization are shown in the right panel. (B) Phenotypes of the parental strains under control conditions and under oxidative stress. (C) Significant correlations between growth in the control condition and growth under oxidative stress at 8 dpi. (D) Significant correlation between relative growth and growth under control conditions at 8 dpi.

Melanization also decreased significantly (relative gray value > 1) under oxidative stress at 8 dpi in both segregating populations (Figure 1A, middle panel), though four offspring from the 3D1 × 3D7 cross showed increased melanization under oxidative stress. At 12 dpi, 36 progeny from the 1E4 × 1A5 cross and 62 progeny from the 3D1 × 3D7 cross became more melanized under oxidative stress (relative gray value < 1), indicating that oxidative stress induced more melanization for those strains at a later stage of growth. To assess how melanin production correlates with colony growth under oxidative stress, simple linear regressions were conducted between relative growth and relative melanization for both time points. A significant negative correlation (p < 0.01) was found between relative gray value and relative growth in both crosses at 8 dpi (Figure 1A, right panel), indicating that a higher reduction in melanization is associated with a higher inhibition in growth under oxidative stress. However, significant correlations between the absolute growth and absolute melanization were also found at 8 dpi for both crosses under both oxidative stress and control conditions (Figure S1), indicating that melanization is associated with growth independent of oxidative stress. In addition, for both crosses under oxidative stress at 12 dpi, the strains with an increase in melanization did not consistently show an increase in growth. On the contrary, most of those strains showed decreased growth under oxidative stress (Figure 1A, right panel). These results suggest that melanization is mostly independent of oxidative stress.

All traits exhibited high values in broad-sense heritability (0.98 – 0.56, Table S1), indicating that the phenotypic variation among offspring was mainly due to genetic variation. The transgressive segregants for these traits likely resulted from recombination of different alleles among several different genes affecting each trait.

### Genetic architecture of growth and melanization under oxidative stress

To investigate the genetic architectures of growth and melanization in response to oxidative stress, interval mapping using a single-QTL genome scan was performed in the two segregating populations. The permutation analysis for all genome scans resulted in threshold LOD scores ranging between 3.4-3.7. For the 1A5 × 1E4 cross, genome scans for the colony radius under oxidative stress (orange line) identified four significant QTLs on chromosomes 1, 3, 8 and 12 at 8 dpi, three of which were also found at 12 dpi (Figure 2A). QTLs on chromosomes 1 and 12 were not found in the genome scan for colony radius in the control environment (blue line), suggesting that these QTLs are specific for growth under oxidative stress. The chromosome 8 QTL for colony radius under oxidative stress had a much higher LOD score and a narrower confidence interval than the chromosome 8 QTL in the control environment at both time points (Figure 2A), suggesting that the chromosome 8 QTL under oxidative stress may be specifically associated with the oxidative stress environment. Multiple peaks are visible in the LOD plots of the chromosome 8 QTL under control conditions (blue line), which may contribute to the wide confidence interval (127 – 430 cM) of this QTL. Genome scans for relative colony radius (yellow line) identified two QTLs on chromosome 1 and 8 at 8 dpi, the same QTLs that were identified for colony radius under oxidative stress (Figure 2A). The chromosome 8 QTL for colony radius under oxidative stress at 8 dpi explained 26% of the phenotypic variation (Table 1A) and resulted in the narrowest 95% confidence interval at this time point (203 – 218 cM), which contained only 36 genes (Table 2).

**Table 1.** Summary of the non-overlapping QTLs with the narrowest confidence intervals associated with colony growth and melanization under oxidative stress.

| Cross | Trait | Chr | Highest LOD | Dpi | Var. (%) | CI (bp) | No. genes | Possible candidates | Control |
| --- | --- | --- | --- | --- | --- | --- | --- | --- | --- |
| 1E4 x 1A5 | Colony Radius | 1 | 4.49 | 8 | 10.04 | 1917295-2665777 | 284 |  | No |
|  | Relative Melanization | 1 | 3.97 | 8 | 4.96 | 262715-733242 | 136 |  | Yes |
|  | Relative Melanization | 2 | 11.11 | 8 | 15.77 | 1589560-1793667 | 67 |  | Yes |
|  | Colony Radius | 3 | 6.38 | 8 | 5.22 | 465107-1344743 | 231 | Cu/Zn SOD, Fe/Mg<br>SOD | Yes |
|  | Melanization | 6 | 3.67 | 12 | 5.84 | 1041770-1583118 | 165 |  | No |
|  | Colony Radius | 8 | 15.82 | 8 | 24.90 | 496559-578252 | 35 |  | Yes |
|  | Relative Melanization | 8 | 7.45 | 8 | 9.51 | 114401-346112 | 78 | Glucose-6-phosphate 1-dehydrogenase (*zwf1*) | Yes |
|  | Relative Growth Rate | 12 | 5.74 | 8-12 | 4.42 | 1110580-1334901 | 82 | Tyrosine protein<br>phosphatase (*yvh1*) | No |

B
| Cross | Trait | Chr | Highest LOD | Dpi | Var. (%) | CI (bp) | No. genes | Possible candidates | Control |
| --- | --- | --- | --- | --- | --- | --- | --- | --- | --- |
| 3D1 x 3D7 | Colony Radius | 1 | 5.59 | 8 | 8.96 | 1246555-2072715 | 262 | Catalase-peroxidase (*ZtCpx2*) | No |
|  | Relative Radius | 1 | 4.86 | 8 | 8.09 | 5256728-5576484 | 104 | Glutaredoxin | No |
|  | Colony Radius | 3 | 5.63 | 12 | 6.53 | 2983747-3365135 | 135 | Catalase (*ZtCat1*) | Yes |
|  | Relative Radius | 7 | 3.68 | 8 | 4.83 | 1363626-1930081 | 201 |  | No |
|  | Melanization | 7 | 4.31 | 8 | 7.20 | 529943-652007 | 50 | Trehalose-phosphatase | No |
|  | Colony Radius | 8 | 3.83 | 12 | 9.72 | 843926-1221280 | 157 | Catalase-peroxidase (*ZtCpx1*) | Yes |
|  | Relative Radius | 10 | 14.85 | 8 | 21.06 | 749521-758289 | 5 |  | Yes |
|  | Relative Growth Rate | 10 | 4.53 | 8-12 | 7.30 | 804952-1126067 | 124 |  | No |
|  | Melanization | 11 | 9.82 | 12 | 14.33 | 514048-879885 | 126 | *PKS1, Zmr1* | Yes |

Chr = chromosome, Dpi = Days post inoculation, Var. = variation explained by the QTL, CI = Confidence interval, Control = Overlaps with QTLs found in the control environment.

**Figure 2.**
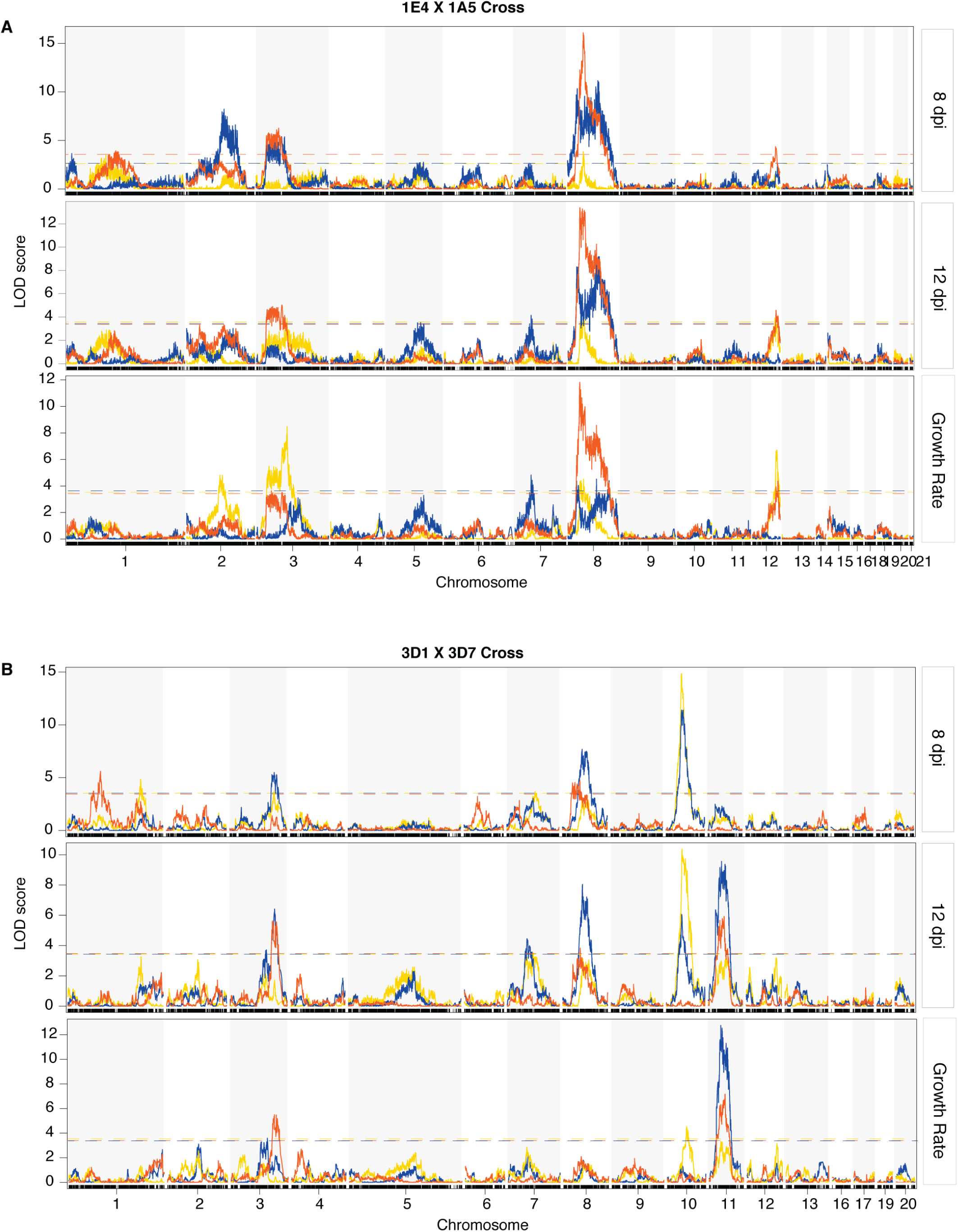
Genetic architecture of growth-related traits. (A) Interval mapping for the 1E4 × 1A5 cross. (B) Interval mapping for the 3D1 × 3D7 cross. The blue line shows the mapping results for colony radius under control conditions, the orange line shows the mapping results for colony radius under oxidative stress and the yellow line shows the mapping results for relative growth. The vertical axis shows the log10 likelihood ratio (LOD score), and the horizontal axis indicates the chromosome number. The horizontal dashed lines in the figure indicate the LOD threshold estimated from 1000 permutations of the genome-wide scan.

**Table 2.** Genes found within the QTL confidence interval on chromosome 8 for growth under oxidative stress at 8 dpi in the 1E4 × 1A5 cross.

| 1A5<br>Gene ID | IPO323<br>Gene ID | IPR_description | Protein polymorphism |  | Homology to *Saccharomyces cerevisiae* |  |  |  |  |
| --- | --- | --- | --- | --- | --- | --- | --- | --- | --- |
|  |  |  | Percentage | Amino<br>Acid | Gene | Coverage | E value | Identity | Description |
| 1A5.g8236 | NA | Lipase, GDXG,<br>putative serine<br>active site,<br>Alpha/Beta<br>hydrolase fold | <1% | 3 |  |  |  |  |  |
| 1A5.g8237 | 8_00172 | Zinc finger,<br>MYND-type | - | very<br>polymor-<br>-phic |  |  |  |  |  |
| 1A5.g8238 | 8_00173 | Arginine<br>biosynthesis<br>protein ArgJ | 2% | 10 | Arg7p | 93% | 5.00E-132 | 45.81% | Ornithine acetyltransferase,<br>involved in ornithine biosynthesis. |
| 1A5.g8239 | 8_00174 | Peptidase M24,<br>Creatinase | 2% | 11 | Fra1p | 99% | 0 | 44.02% | Involved in the regulation of the<br>iron regulon in responses to<br>decreased mitochondrial iron-<br>sulfur cluster synthesis. |
| 1A5.g8240 | 8_00175 | Catechol<br>dioxygenase,<br>Intradiol ring-<br>cleavage<br>dioxygenase | <1% | 2 |  |  |  |  |  |
| 1A5.g8241 | 8_00176 | NA | - | Another<br>298 aa<br>more in<br>1E4 |  |  |  |  |  |
| 1A5.g8242 | 8_00177 | NADPH-dependent FMN reductase-like, Arsenate resistance ArsH, Flavoprotein-like domain | 0% |  |  |  |  |  |  |
| 1A5.g8243 | 8_00178 | Zinc finger, double-stranded RNA binding, Splicing factor 3A subunit 3, C2H2-type | <1% | 1 | Prp9p | 76% | 3.00E-49 | 30.05% | Necessary for binding of the U2 snRNP to the pre-mRNA in an early step of spliceosome assembly. |
| 1A5.g8244 | 8_00179 | ATP-dependent RNA helicase DEAD-box | 0% |  | Fal1p | 98% | 0 | 64.30% | Involved in 40S ribosomal subunit biogenesis. |
| 1A5.g8245 | 8_00180 | Peptidyl-tRNA hydrolase | 6% | 14 (+2 indels) | Pth1p | 64% | 9.00E-12 | 34.68% |  |
| 1A5.g8246 | 8_00181 | Winged helix-turn-helix DNA-binding domain | 0% |  |  |  |  |  |  |
| 1A5.g8247 | 8_00182 | TB2/DP1/HVA22-related protein | 0% | 0 | Yop1p | 90% | 3.00E-34 | 39.63% | Involved in membrane/vesicle trafficking. |
| 1A5.g8248 | 8_00183 | SNF2-related, P-loop containing nucleoside triphosphate hydrolase, Sterile alpha motif/pointed domain, Helicase superfamily 1/2, ATP-binding domain | 1% | 3 |  |  |  |  |  |
| 1A5.g8249 | 8_00184 | Glutathione synthase | 1% | 2 | Gsh2p | 98% | 1.00E-122 | 40.97% | Involved in step 2 of the biosynthesis of glutathione. |
| 1A5.g8250 | 8_00185 | Domain of unknown function DUF4203 | 3% | 34 |  |  |  |  |  |
| 1A5.g8251 | 8_00186 | Uncharacterized protein family UPF0642 | <1% | 1 |  |  |  |  |  |
| 1A5.g8252 | 8_00187 | Zinc finger, MIZ-type, E3 SUMO protein ligase, SAP domain, PINIT domain | 0% | 0 |  |  |  |  |  |
| 1A5.g8253 | 8_00188 | DNA repair protein, Swi5 | 7% | 13 |  |  |  |  |  |
| 1A5.g8254 | 8_00189 | Domain of unknown function DUF2427 | <1% | 3 | Ytp1p | 62% | 3.00E-69 | 41.69% | Can function as a molecular chaperone, and controls proteins involved in metabolism, stress response, and DNA regulation. |
| 1A5.g8255 | 8_00190 | Zinc finger, MYND-type | <1% | 3 |  |  |  |  |  |
| 1A5.g8256 | 8_00191 | Zinc finger, MYND-type | 3% | 6 |  |  |  |  |  |
| 1A5.g8257 | 8_00192 | Zinc finger, MYND-type | <1% | 1 |  |  |  |  |  |
| 1A5.g8258 | 8_00193 | Zinc finger, MYND-type | 17% | 31 (+4 indels) |  |  |  |  |  |
| 1A5.g8259 | 8_00194 | Zinc finger, MYND-type | 5% | 10 |  |  |  |  |  |
| 1A5.g8260 | 8_00195 | - | >50% | very polymorphic |  |  |  |  |  |
| 1A5.g8261 | NA | Zinc finger, MYND-type | >50% | very polymorphic |  |  |  |  |  |
| 1A5.g8262 | NA | Zinc finger, MYND-type | <1% | 2 |  |  |  |  |  |
| 1A5.g8263 | NA | - | >50% | very polymorphic |  |  |  |  |  |
| 1A5.g8264 | NA | Zinc finger, MYND-type | 0% | 0 |  |  |  |  |  |
| 1A5.g8265 | 8_00199 | Zinc finger, MYND-type | <1% | 2 |  |  |  |  |  |
| 1A5.g8266 | 8_00200 | Sec8 exocyst complex component specific domain | 0% | 0 | Sec8p | 89% | 4.00E-55 | 22.26% | Involved in the docking of exocytic vesicles with fusion sites on the plasma membrane. Some alleles of this gene are temperature sensitive. |
| 1A5.g8267 | 8_00201 | Tetratricopeptide repeat | <1% | 1 | Ctr9p | 90% | 1.00E-102 | 27.56% | Involved in transcription initiation and elongation. Also has a role in transcription-coupled histone modification and chromosome segregation. |
| 1A5.g8268 | 8_00202 | Zn(2)-C6 fungal-type DNA-binding domain | <1% | 4 |  |  |  |  |  |
| 1A5.g8269 | 8_00203 | Pyruvate kinase-like, Molybdenum cofactor sulfurase, C-terminal, N-terminal beta barrel | <1% | 1 |  |  |  |  |  |
| 1A5.g8270 | NA | - | <1% | 1 |  |  |  |  |  |
| 1A5.g8271 | 8_00204 | Zn(2)-C6 fungal-type DNA-binding domain | <1% | 4 | Upc2<br>p | 82% | 1.00E-11 | 24.52% | Involved in activation of anaerobic genes, sterol uptake and regulation of the sterol biosynthesis. |

Note: protein sequences were compared between the 1A5 and 1E4 strains. Polymorphism $\leq 1\%$ appear as 1%. The amino acid substitutions were not counted for genes with polymorphism $>50\%$ . Aa = amino acids, indels = insertions and deletions.

For the 3D7 × 3D1 cross, colony radius under oxidative stress at 8 dpi mapped to two QTLs located on chromosomes 1 and 8, while colony radius at 12 dpi mapped to two QTLs on chromosomes 3 and 11 and a QTL on chromosome 8 that overlapped with the chromosome 8 QTL at 8 dpi (Figure 2B). The chromosome 1 QTL did not overlap with QTLs found under control conditions, suggesting it plays a role in oxidative stress tolerance. Relative growth mapped to two weak QTLs on chromosomes 1 and 7, and a large QTL (LOD = 15) on chromosome 10 at both time points (Figure 2B, top and middle panels). The chromosome 10 QTL overlapped with the QTL in the control environment but with a higher LOD score and narrower confidence intervals at both time points, suggesting a higher significance of association for relative growth than for growth in the control environment. This chromosome 10 QTL accounted for 23% and 16% of the phenotypic variation in relative growth at 8 and 12 dpi, respectively. The narrowest confidence interval for this chromosome 10 QTL was found at 8 dpi, covering an 8751 bp region that included only 5 genes (Table 1B).

The genome scan for melanization under oxidative stress in the 1A5 × 1E4 cross identified four QTLs on chromosomes 2, 3, 8 and 12 at 8 dpi (Figure 3A). All these QTLs, except the chromosome 8 QTL, overlapped with the QTLs found for melanization in the control environment (Table 1A). Relative melanization at 8 dpi mapped to the same QTLs found under control conditions, though the LOD scores were lower (Figure 3A). The relative melanization rate mapped to a large chromosome 3 QTL, the same QTL that was found for melanization under oxidative stress at 12 dpi and in the control environment at 8 dpi. The chromosome 8 QTL for melanization under oxidative stress at 8 dpi was the only oxidative stress-specific QTL. This QTL overlapped with the QTL found for growth under oxidative stress at 8 dpi.

**Figure 3.**
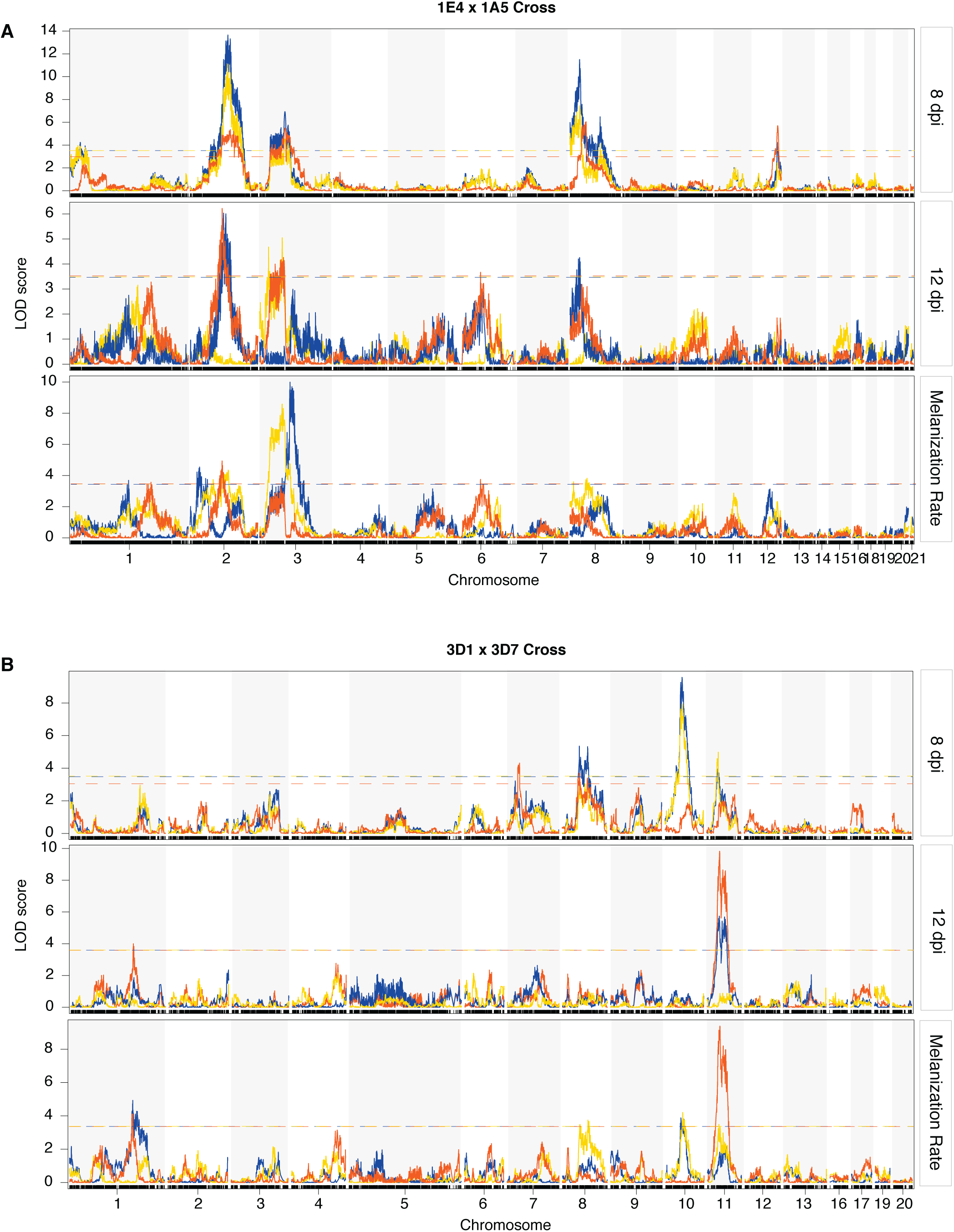
Genetic architecture of melanization-related traits. (A) Interval mapping for the 1E4 × 1A5 cross. (B) Interval mapping for the 3D1 × 3D7 cross. The blue line presents the mapping results for colony melanization under control conditions, the orange line presents the mapping results for colony melanization under oxidative stress and the yellow line presents the mapping results for relative melanization. The vertical axis shows the log10 likelihood ratio (LOD score), and the horizontal axis indicates the chromosome number. The horizontal dashed lines in the figure indicate the LOD threshold estimated from 1000 permutations of the genome-wide scan.

In the 3D1 × 3D7 cross, colony melanization under oxidative stress at 8 dpi mapped to QTLs on chromosomes 7 and 8 and relative melanization mapped to QTLs on chromosomes 10 and 11. Only the chromosome 7 QTL did not overlap with QTLs found in the control environment (Figure 3B). Colony melanization at 12 dpi mapped to QTLs on chromosomes 1 and 11. The chromosome 1 QTL did not overlap with the control environment. The chromosome 11 QTLs at 12 dpi were much stronger under oxidative stress than in the control environment. The chromosome 11 QTLs included the *PKS1* and *Zmr1* genes that regulate the production of melanin in *Z. tritici* (Lendenmann *et al*. 2014; Krishnan *et al*. 2018).

To summarize, QTLs specifically associated with growth under oxidative stress were found on chromosomes 1 and 12 in the 1E4 × 1A5 cross and on chromosomes 1 and 7 in the 3D1 × 3D7 cross. The chromosome 8 QTL in the 1E4 × 1A5 cross covered a narrower interval and had a higher LOD score for growth under oxidative stress than for growth under control conditions, suggesting it may affect growth under oxidative stress. Similarly, the chromosome 10 QTL in the 3D1 × 3D7 cross had a higher LOD score for relative growth than for absolute growth in the control environment, suggesting it may also affect growth under oxidative stress. QTLs found for melanization under oxidative stress largely overlapped with the QTLs found in the control environment, suggesting that these QTLs were not specifically associated with oxidative stress and/or that they showed pleiotropic effects. All unique QTLs (i.e. lacking overlapping confidence intervals) found under oxidative stress conditions were summarized in Table 1, which shows only the QTLs with the narrowest confidence interval in cases where there were overlapping QTLs. Detailed information summarizing all the significant QTLs for each trait are presented in Tables S2 and S3.

### Identification of candidate genes within QTL confidence intervals

Only QTLs associated with tolerance to oxidative stress that had a LOD > 10 and confidence intervals containing less than 40 genes were considered for identification of candidate genes. The chromosome 8 QTL for growth under oxidative stress in the 1E4 × 1A5 cross, the chromosome 10 QTL for relative growth and the chromosome 11 QTL for melanization in the 3D1 × 3D7 cross satisfied these criteria. Because the confidence interval of the chromosome 11 QTL for melanization in the 3D1 × 3D7 cross overlapped with the previously identified QTL for melanization in the same cross (Lendenmann *et al*. 2014), we consider this chromosome 11 QTL to be the same QTL reported previously. The candidate genes for this chromosome 11 QTL were already functionally characterized (Krishnan *et al*. 2018), leading to discovery of polymorphisms affecting transcription of *Zmr1* that explained the observed differences in melanization. As we believe that the same polymorphisms can explain the QTL discovered in this independent experiment, we will not discuss this QTL further.

The chromosome 8 QTL (LOD = 15.8) for colony radius under oxidative stress at 8 dpi in the 1A5 × 1E4 cross (Figure 4) contains 36 genes. The marker with the highest LOD is located in a gene 1A5.g8250 (Zt09_8_00185) without a known functional domain (Figure 4C). BLAST analyses to the NCBI database of protein sequences identified eight genes with high similarity to characterized genes in yeast (taxid:4932) (Table 2). Among these genes, glutathione synthetase (1A5.g8249) and Fra1 (1A5.g8239) could potentially affect growth under oxidative stress, according to studies conducted with *Saccharomyces cerevisiae* (Grant *et al*. 1997; Kumánovics *et al*. 2008). Many genes found in this confidence interval have no homology to characterized genes, but are likely to be involved in the cell cycle and/or basic metabolism based on the presence of functional domains associated with DNA repair (1A5.g8253), zinc fingers (1A5.g8255 - 8265) and RNA polymerase (1A5.g8267). A high degree of DNA polymorphisms between the two parental strains, including a transposable element insertion, were found in this confidence interval (Figure 4D). These polymorphisms could affect the expression of genes, protein structures or protein maturation processes.

**Figure 4.**
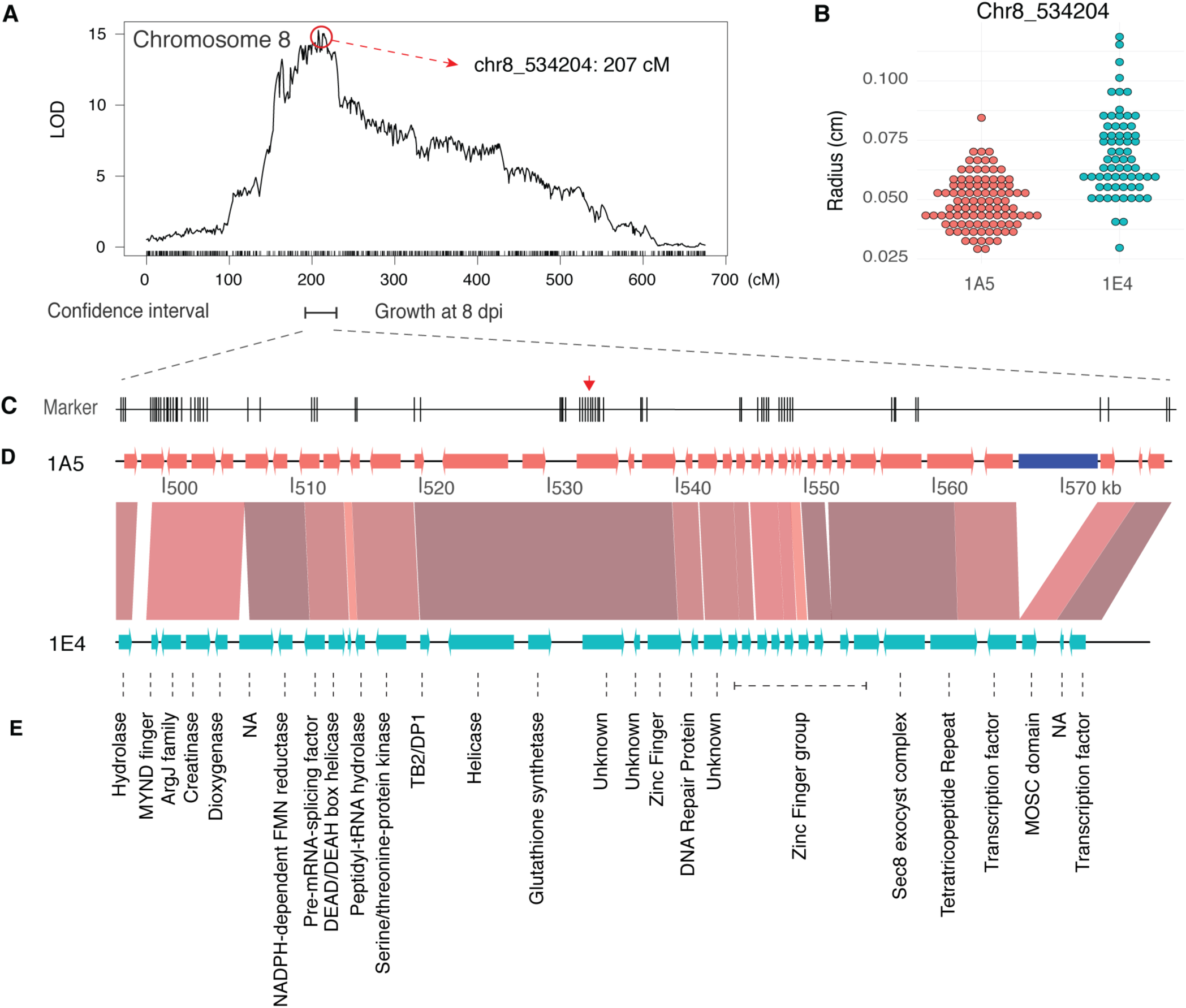
The chromosome 8 QTL for growth under oxidative stress in the 1E4 × 1A5 cross. (A) LOD plot of the chromosome 8 QTL for growth under oxidative stress at 8 dpi. The y axis indicates the LOD score, and the × axis indicates the genetic distance (centiMorgans) along the chromosome. The red arrow indicates the marker with the highest LOD, which resides at 534204 bp for growth at 8 dpi. The 95% Bayesian confidence interval is shown below the LOD plot. (B) Allele effect of the chromosome 8 QTL: individuals with the 1A5 allele at 534204 bp have a smaller colony radius under oxidative stress than individuals with the 1E4 allele at 534204 bp. Each closed circle represents a different offspring. (C) SNP markers in the confidence intervals are shown as vertical lines. The red arrow indicates the location of the marker with the highest LOD. (D) Synteny plot showing the DNA polymorphisms in this genomic region. The red segments indicate regions with sequence identity greater than 90%. The darker the red color, the lower the degree of polymorphism in the genomic region. The arrows represent genes, and the blue block represents a transposable element. (E) Functional domains in the 36 genes within the confidence interval. Details for these genes are presented in Table 2.

The QTL on chromosome 10 for relative growth at 8 dpi in the 3D1 × 3D7 cross contained only five genes (Figure 5E), including a ferric reductase (3D7.g9787), a major facilitator superfamily transporter (MFS, 3D7.g9788), an acetyltransferase (3D7.g9789) and two genes (3D7.g9786 and 3D7.g9790) encoding proteins without known functional domains. BLAST analysis of the protein sequence of the ferric reductase showed that this gene contains a NOX_Duox_like_FAD_NADP domain (E value = 2.36e-43) and a ferric reductase domain belonging to the cytochrome b superfamily (E value = 4.83e-25). However, a BLAST analysis between this NOX in *Z. tritici* and NOXs characterized in other fungi such as NOXB (CAP12517) and NOXA (CAP12516) from *Botrytis cinerea* (Segmüller *et al*. 2008) showed low similarity (E value = 5e-07, Identity = 23%, Coverage = 30%), suggesting that this protein represents a previously uncharacterized NOX in fungi. BLAST analysis of the MFS transporter indicated that this gene has a domain similar to a D-galactonate transporter (E value = 3.63e-57), which is associated with transport of proteins and carbohydrates. BLAST analysis of the acetyltransferase indicated that it belongs to the acetyltransferase_10 superfamily, which could affect transcriptional elongation. Many polymorphisms were found for the ferric reductase protein sequences among the two parental strains (3D1 and 3D7), in the 5’-UTR of the MFS gene and in the 3’-UTR of the acetyltransferase (Figure 5D and 5E). Four out of the five genes in this confidence interval showed significant differential expression between the two parental strains in yeast sucrose broth (YSB) media (Figure 5F). Gene 3D7.g9786 and the ferric reductase gene were downregulated under starvation conditions, while the MFS and acyltransferase genes were upregulated under starvation conditions (Figure 5F).

**Figure 5.**
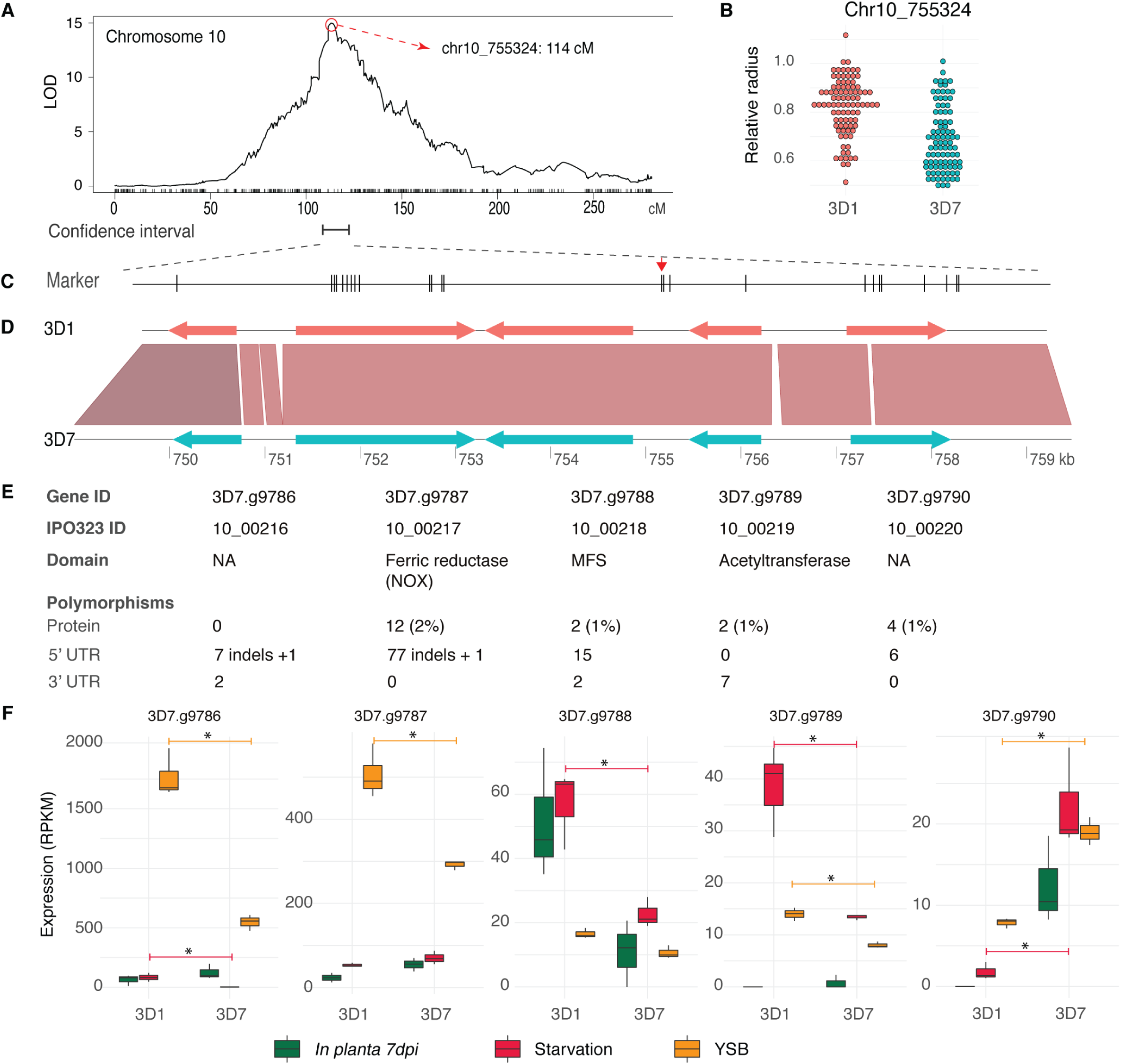
The chromosome 10 QTL for relative growth at 8 dpi in the 3D1 × 3D7 cross. (A) LOD curve of the chromosome 10 QTL. The red arrow indicates the peak marker (chr10_755324). (B) Allele effects at the marker Chr10_755324: individuals with 3D1 allele showed higher average relative growth than individuals with the 3D7 allele. Each closed circle represents an individual. (C) Markers in the confidence interval of this QTL. Red arrow indicates the peak marker. (D) Synteny plot of the confidence interval region between the two parental strains. The darker the red color, the lower the degree of polymorphism in the genomic region. (E) The gene ID in 3D7, the corresponding gene ID in IPO323, the functional domains and the gene polymorphisms associated with (D). Polymorphism numbers indicate the number of substitutions, and indels indicate the number of nucleotides missing in one of the parental strains. The number in brackets indicates the percentage of polymorphic amino acids in the entire protein. (F) Expression values in reads per kilobase of transcript per million mapped reads (RPKM) of the five genes in yeast sucrose broth medium (YSB, control), minimal medium (starvation) and *in planta* infection at 7 dpi. The expression data were collected from three replicates in a previous study (Francisco *et al*. 2018).

### In planta expression profile of the candidate genes

We compared the expression patterns of the candidate genes with known ROS scavengers, including the only known catalase-peroxidases, the only known glutathione peroxidase, four catalases and six SODs in *Z. tritici*. We also included homologs of the two major regulators of oxidative stress, *ZtYap1* (Yang *et al*. 2015) and *MgHog1* (Mehrabi *et al*. 2006), and the characterized homolog *NOXa* in *Z. tritici* (Choi *et al*. 2016). The expression patterns of the candidate genes in the chromosome 8 and 10 QTLs and selected genes related to ROS elimination (Figure S4) separated them into three groups: highly expressed in the biotrophic (7 dpi), necrotrophic (11 and 14 dpi) or saprotrophic (21 dpi) phase, according to the infection time course shown earlier (Palma-Guerrero *et al*. 2016). The DNA repair protein (Swi5) in the chromosome 8 QTL and gene 3D7.g9790 in the chromosome 10 QTL showed their highest expression during the early biotrophic phase (3 dpi). The ferric reductase reached its highest expression at 7 dpi, together with genes such as catalase 1 (*ZtCat1*), *ZtCpx1* and *Yap1*, matching the timing of the first ROS burst reported in a previous study (Shetty *et al*. 2003). The gene 3D7.g9786 (without a functional domain), the MFS and the acetyltransferase in the chromosome 10 QTL showed the highest expression at the early necrotrophic phase (11 dpi), together with catalase-peroxidase 2 and glutathione peroxidase. Glutathione synthetase, one of the most compelling candidate genes for the chromosome 8 QTL, showed the highest expression at 14 dpi, together with its adjacent gene 1A5.g8250 and two SODs, matching the timing of the second ROS burst reported previously (Shetty *et al*. 2003). The SODs found in our QTLs showed their highest expression during the saprotrophic phase, together with catalase 2.

## DISCUSSION

Tolerance to exogenous oxidative stress is important for most plant pathogens, as it is associated with their pathogenicity and viability during host infection. In this study, we investigated the genetic architecture of this trait in two segregating populations of the fungal wheat pathogen *Z. tritici*. The large variation found for both growth and melanization in both control and oxidative stress environments were largely explained by genetic variation. Mapping these traits resulted in sixteen significant QTLs without overlapping confidence intervals among the two crosses. Among these 16 QTLs, four were specifically associated with the oxidative stress environment. Two other QTLs with high LOD scores were significantly associated with growth under oxidative stress or sensitivity to oxidative stress. A detailed examination of these two QTLs led to identification of candidate genes that may affect oxidative stress tolerance of *Z. tritici*.

### Reproducibility of results from independent QTL analyses

In previous QTL mapping investigations oriented around temperature sensitivity (Lendenmann *et al*. 2016), melanization (Lendenmann *et al*. 2014) and fungicide tolerance (Lendenmann *et al*. 2015), the experiments were carried out using PDA media either amended or without fungicides with growth at either 15 or 22 °C. Our study used the same fungal strains and the same culture media purchased from the same company, but with growth at 18 °C. We measured growth and melanization using image analysis as presented in the earlier studies. Hence, we expected to reproduce some of the QTL peaks involved in general regulation of growth and melanization in our study.

Four of the QTLs identified in this study were found in similar genomic regions in the earlier studies. The chromosome 8 QTL found for growth in the control environment in the 1E4 × 1A5 cross was also found when mapping growth at 15 °C (Lendenmann *et al*. 2016). The confidence interval of this QTL overlapped with the confidence intervals of QTLs found for growth at 22 °C (this confidence interval included genes 1A5.g8145-8189) and fungicide sensitivity (this confidence interval included genes 1A5.g8172-8197) (Lendenmann *et al*. 2015; 2016). The chromosome 10 QTL for relative growth under oxidative stress in the 3D1 × 3D7 cross was also found at a similar position (including genes 3D7.g9787-9790 in this study and genes 3D7.g9793-9801 in the previous study) when mapping for relative growth rate at 15 and 22 °C (Lendenmann *et al*. 2016). Results of both studies indicated a higher LOD of the chromosome 10 QTL for the relative trait (sensitivity) than for the control environment. Re-mapping of the thermal sensitivity trait of Lendenmann *et al*. 2016 (which used IPO323 as the reference genome), using our new genetic map (using 3D7 as the reference genome), resulted in the same confidence interval for the chromosome 10 QTL (including genes 3D7.g9787-9789) affecting thermal sensitivity as for the oxidative stress sensitivity trait considered in this study. The chromosome 2 QTL for melanization in the 1E4 × 1A5 cross was also found in the previous mapping for melanization (Lendenmann *et al*. 2014), with the confidence intervals in the same region (including genes 1A5.g2705-2769 in this study and genes 1A5.g2708-2760 in the previous study). The chromosome 11 QTL found in this study for melanization (including genes 3D7.g10272-10363) overlapped with the chromosome 11 QTL for melanization found in a previous study, which led to the discovery that the *Zmr1* gene was regulating the production of melanin (Lendenmann et al 2014; Krishnan *et al*. 2018).

The reproducibility of QTL peaks between independent studies conducted under similar experimental conditions illustrates the robustness and accuracy of our phenotyping methods. The overlap of growth QTLs under different environmental conditions suggests that these QTLs contain genes that generally regulate growth under many different conditions. But the overlap between QTLs affecting melanization and QTLs affecting growth suggests that there is a connection between these two traits, indicative of pleiotropy and possible tradeoffs. The same pattern was found in earlier studies (Lendenmann et al 2015).

### Role of melanization under oxidative stress

Though melanin was suggested to act as a physiological redox buffer that helps protect fungal cells against ROS (Jacobson 2000), to our knowledge no direct empirical evidence was found to support this assumption in fungi. In our study, we compared colony melanization under oxidative stress with a control environment and found stress-induced melanization in some strains (Figure 1A, middle panel). A significant positive correlation was found between decreased melanization (relative gray value < 1) and inhibited growth (relative growth > 1) at 8 dpi in both populations (Figure 1A, right panel). This correlation appears to support the hypothesis that melanin increases tolerance to oxidative stress. However, this correlation could simply reflect the correlation between the absolute melanization and absolute growth at 8 dpi in both environments (Figure S1), as the correlations between the relative values were only significant when the correlations between the absolute values were significant. Our finding of overlapping QTLs for relative melanization and melanization in the control environment at 8 dpi (Figure 3A) supports this explanation. We did not find a consistent correlation between increased melanization and increased growth under oxidative stress among the thirty-seven strains from the 1E4 × 1A5 cross that showed ROS-induced melanization (relative gray value < 1) at 12 dpi (Figure 1A, middle panel). This result also suggests that the production of melanin does not contribute to higher oxidative stress tolerance.

### Growth under oxidative stress

We found weak but significant correlations between growth under oxidative stress and growth in the control environment for both populations (Figure 1C). The highest correlation (R^2^ = 0.25) between these two traits was found in the 1E4 × 1A5 cross at 8 dpi. Closer examination of sixteen strains from this cross exhibiting extremely high (the eight highest) or low (the eight lowest) growth values under oxidative stress showed that the biggest difference in colony size between the two groups of strains was exhibited only under oxidative stress (Figure S2). In addition, differences in colony size in the control environment were also found between the two groups of the strains shown in Figure S2. These results suggest that some of the genes controlling growth under oxidative stress and controlling growth in a non-stressful environment are linked or are the same. Overlapping QTLs for growth in the control environment and under oxidative stress were found on chromosomes 3 and 8 in both crosses (Figure 2A). This suggests that both of these chromosome regions carry genes that may play a general role in growth *in vitro*.

In the confidence interval of the chromosome 8 QTL under oxidative stress, we found many genes encoding domains associated with oxidative stress responses as well as general growth (Figure 4D, Table 2). Genes encoding glutathione synthetase, DNA repair protein, zinc fingers and transcription factors were found in this confidence interval. Focusing on the genes in this QTL that are mostly likely to affect oxidative stress tolerance, we considered gene 1A5.g8249 (similar to the glutathione synthetase gene (*Gsh2*) in *S. cerevisiae*) as one of our top candidates because: (1) Glutathione synthetase was found to affect hydrogen peroxide tolerance in *S. cerevisiae* (Grant *et al*. 1997), *Candida glabrata* (Gutiérrez-Escobedo *et al*. 2013) and *Aspergillus nidulans* (Bakti *et al*. 2017), with the depletion of this enzyme resulting in slower growth in H_2_O_2_. (2) Glutathione plays an essential role in maintaining the cellular redox environment (Aquilano *et al*. 2014; Corso & Acco 2018), which is important for fungal cell development (Samalova *et al*. 2013). Glutathione synthetase is one of only two enzymes involved in the synthesis of glutathione and *Z. tritici* has only one pair of these enzymes in its genome. (3) The glutathione synthetase gene is very close to the peak marker (Figure 4C) and possessed two amino acid polymorphisms in its protein sequence between the two parental strains which could contribute to the observed phenotypic variation. It is also noteworthy that the adjacent gene (1A5.g8250) has 34 amino acid polymorphisms in its protein sequence and that the peak marker resides in this gene. So gene 1A5.g8250 is also a strong candidate for this QTL. But we cannot exclude the possibility that other genes in this QTL encode proteins affecting the oxidative stress response and/or cell growth, such as a peptidase gene (1A5.g8239) involved in iron regulon regulation, a gene with a DNA repair protein domain (1A5.g8253) and a group of zinc finger proteins likely affecting protein interactions. It is also possible that different genes in this confidence interval independently contribute to growth under oxidative stress and growth under non-stressed conditions. Whether it is one gene or several genes in this confidence interval contributing to the variation in growth under oxidative stress will be revealed by future functional studies.

The QTLs on chromosome 1 and chromosome 12 for the 1E4 × 1A5 cross and the QTLs on chromosomes 1 and 7 for the 3D1 × 3D7 cross were specifically associated with oxidative stress. These QTLs encompass homologs of genes known to be involved in oxidative stress tolerance including glutamate-cysteine ligase (3D7.g1830), which is the other enzyme involved in the biosynthesis of glutathione and known to be essential for growth under oxidative stress (Grant *et al*. 1997; Bakti *et al*. 2017). Catalase-peroxidase 2 (*ZtCpx2*, 3D7.g1854) was found to be important in tolerance to high levels of oxidative stress in *Z. tritici* in a previous study (Gohari A. M. 2015) (Table 1). A large number of genes with unknown functions were also found in these oxidative stress specific QTLs, providing many opportunities to identify new candidate genes related to oxidative stress tolerance. Interestingly, other known ROS regulators such as catalase 1 (*ZtCat1*, 3D7.g4375) and superoxide dismutase (Cu/Zn SOD 1A5.g3638 and Fe/Mg SOD 1A5.g3652) were found in QTLs that overlapped with QTLs found in the control environment (Table 1). Given the small effects and large confidence intervals of these QTLs, together with the lack of protein polymorphism in the associated *SOD*s and *ZtCat1*, it is difficult to ascertain whether these genes are involved in the growth under oxidative stress or in non-stressed conditions. Functional studies using these well-known ROS regulators will help to illustrate their effects on oxidative stress tolerance in *Z. tritici*.

### Tradeoff between growth and growth inhibition under oxidative stress

The phenotypic data showed a negative correlation between growth inhibition under oxidative stress and growth in the control environment, indicating that strains growing the fastest in the control environment are the most sensitive to oxidative stress. This correlation was strongest in the 3D1 × 3D7 cross at 8 dpi (R^2^ = 0.59) (Figure 1D), suggesting that the large variation in growth inhibition may result from variation in growth under non-stressful conditions. To illustrate this correlation, colony images from the eight segregating strains with the highest relative growth value and eight strains with the lowest relative growth were compared (Figure S3). Bigger differences in colony growth were found in the control environment compared to the oxidative stress environment, supporting our hypothesis that the difference in relative growth largely reflects growth differences under non-stressful conditions. Mapping growth inhibition (measured as relative growth) in the 3D1 × 3D7 cross further supported this hypothesis, as a single QTL (LOD = 15) on chromosome 10 overlapped with the chromosome 10 QTL (LOD = 11) found for growth in the control environment (Figure 2B). Because a previous study found that the same QTL was associated with temperature sensitivity (Lendenmann *et al*. 2016), we postulate that this QTL may include genes that affect sensitivity to several stresses.

Under this hypothesis, we sought candidate genes in the chromosome 10 QTL associated with both fungal growth and stress sensitivity. Among the five genes in the confidence interval (Figure 4C), the ferric reductase with a NOX domain (3D7.g9789) has a function directly related to fungal growth. NOX has been shown to be involved in many fungal developmental processes including cell differentiation, hyphal tip growth and spore germination (Egan *et al*. 2007; Segmüller *et al*. 2008; Ryder *et al*. 2013; Samalova *et al*. 2013). These functions are likely to be highly associated with the colony growth measured in this study. NOX was also reported to be involved in the oxidative stress response in the fungal plant pathogens *Botrytis cinerea* and *Alternaria alternata*, where knocking down either NOXA or NOXB resulted in smaller colonies under oxidative stress. A BLAST analysis of the protein sequence of NOXA from *B. cinerea* resulted in gene 3D7.g686 (NOXa) on chromosome 1, which has been characterized as a homolog of NOX1 from *M. oryzae* and was found to affect virulence in *Z. tritici* (Choi *et al*. 2016). A BLAST analysis of 3D7.g9789 against other fungal genomes resulted in uncharacterized genes, indicating that this NOX (3D7.g9789) is likely an uncharacterized NOX in fungi with an unknown function. As the 3D7 allele is associated with a higher stress sensitivity than the 3D1 allele (Figure 5B), it is possible that the higher sensitivity of the 3D7 allele reflects differences in the protein sequence of the candidate gene between the two parental strains (Figure 5E). It is also possible that the different phenotypes reflect differential expression of the candidate gene as a result of sequence polymorphisms found in the UTR region (Figure 5E). The RNA expression data of the five candidate genes collected from a previously published study of *Z. tritici* (Francisco *et al*. 2019) showed significantly higher expression of the 3D7 allele than the 3D1 allele under both control (YSB) and stress conditions (Figure 5F).

### Complex *in planta* expression of the candidate genes

Plants infected by *Z. tritici* produce ROS during the biotrophic phase (5-7 dpi) and the necrotrophic phase (from around 13 dpi) of infection (Shetty *et al*. 2003). In our study, we found a correlation between the timing of the first ROS burst and the peak of expression (7 dpi) of genes such as *ZtCat1*, *ZtCpx1* and NOX (3D7.g9787), and a correlation between the second burst of ROS and the peak of expression (14 dpi) of genes such as SODs and the glutathione synthetase (Figure S4). These results suggest their possible roles in providing tolerance to exogenous ROS from plant defense around 7 dpi and providing tolerance to endogenous ROS that may peak during fungal sporulation and exogenous ROS released during the collapse of the plant cells around 14 dpi. Interestingly, *ZtCpx2* is only expressed during the transition from the asymptomatic to the necrotrophic phase (11 dpi), which may be correlated with a need to eliminate the large amount of ROS released from dying plant cells (Figure S4). However, the ROS measurements reported in Shetty et al (2003) were performed with a different pathogen strain and host cultivar, which may result in a different timing of this ROS burst. To better understand the roles played by these candidate genes in tolerance to oxidative stress, future studies should consider the effect of different doses of ROS on *Z. tritici* growth, and conduct additional ROS measurements *in planta*.

## CONCLUSION

In this study, we identified genomic regions associated with variation in oxidative stress tolerance using QTL mapping in two crosses of *Z. tritici*. We found QTLs specific for oxidative stress tolerance, and identified candidate genes that may explain a significant portion of the observed variation. Based on our findings, we hypothesize that genes related to fungal cell growth could also contribute to the variation in oxidative stress tolerance.

## ACKNOWLEDGEMENTS

We thank Tiziana Valeria Vonlanthen, Bethan Turnbull, Susanne Dora, Sarah Furler, Alexandra Waltenspühl and Jasmin Wiedmer for helping to conduct experiments. Thanks to Xin Ma and Carolina Francisco for providing the analyzed RNAseq expression data. We are grateful to Andrea Sánchez Vallet and Daniel Croll for critical reading of the manuscript. This research was supported by the Swiss National Science Foundation (31003A_155955 granted to BAM).

